# Morphine-induced hyperalgesia impacts small extracellular vesicle miRNA composition and function

**DOI:** 10.1101/2024.10.17.617815

**Authors:** Deepa Reddy, Zhucheng Lin, Sujay Ramanathan, Xuan Luo, Richa Pande, Yuzhen Tian, Christine Side, Jacqueline M. Barker, Ahmet Sacan, Julie A. Blendy, Seena K. Ajit

**Author notes:** Equal contributions (Deepa Reddy, Zhucheng Lin, Sujay Ramanathan). Corresponding author Seena K. Ajit PhD, Pharmacology & Physiology, Drexel University College of Medicine, 245 North 15th Street, Mail Stop 488, Room 8223, Philadelphia, PA 19102.

## Abstract

Morphine and other synthetic opioids are widely prescribed to treat pain. Prolonged morphine exposure can paradoxically enhance pain sensitivity in humans and nociceptive behavior in rodents. To better understand the molecular mechanisms underlying opioid-induced hyperalgesia, we investigated changes in miRNA composition of small extracellular vesicles (sEVs) from the serum of mice after a morphine treatment paradigm that induces hyperalgesia. We observed significant differential expression of 18 miRNAs in sEVs from morphine-treated mice of both sexes compared to controls. Several of these miRNAs were bioinformatically predicted to regulate cyclic AMP response element binding protein (CREB), a well-characterized transcription factor implicated in pain and drug addiction. We confirmed the binding and repression of *Creb* mRNA by miR-155 and miR-10a. We tested if serum-derived sEVs from morphine-treated mice could elicit nociceptive behavior in naïve recipient mice. Intrathecal injection of 1 μg sEVs did not significantly impact basal mechanical and thermal threshold in naïve recipient mice. However, prophylactic 1 μg sEV administration in recipient mice resulted in faster resolution of complete Freund’s adjuvant-induced mechanical and thermal inflammatory hypersensitivity. Other behaviors assayed following administration of these sEVs were not impacted including sEV conditioned place preference and locomotor sensitization. These results indicate that morphine regulation of serum sEV composition can contribute to analgesia and suggest a potential for sEVs to be a non-opioid therapeutic intervention strategy to treat pain.

## INTRODUCTION

Chronic pain affects more than 100 million people in the United States with the annual cost to society estimated at $635 billion [1]. Morphine and other synthetic opioids are widely prescribed to treat pain, but their continued use can induce a progressive decrease in analgesia (tolerance) and the development of hypersensitivity to painful stimuli or opioid-induced hyperalgesia (OIH) [2,3]. Thus, the treatment meant to relieve pain will paradoxically exacerbate pain. This vicious cycle can lead some patients to develop drug addiction or abuse with negative consequences, severely complicating the management of chronic pain [4,5]. Although multidisciplinary approaches including opioid rotation, opioid tapering, incorporating non-opioid analgesics and non-pharmacological interventions are pursued, OIH remains an unmet clinical need [6] and the mechanisms underlying OIH are not well understood.

Secreted membrane-enclosed vesicles, collectively called extracellular vesicles (EVs), include exosomes, microvesicles, apoptotic bodies, and other EV subsets [7]. Small extracellular vesicles (sEVs), predominantly exosomes, have been implicated for their role in mediating intercellular communication. sEVs are 30–150 nm vesicles that transport biomolecules, including miRNAs, mRNAs, proteins, and lipids in bodily fluids [7,8]. sEVs do not incorporate at random or everything present in the parental cell cytoplasm, suggesting that their synthesis is a well-regulated process that can be dynamically altered by signaling cues [9]. sEVs can cross the blood-brain barrier and have gained significant attention in recent years for their biomarker potential [10,11]. They can mediate communication in the nervous system, which may supplement the known mechanisms of anterograde and retrograde signaling across synapses [12]. While sEVs can have a detrimental role in neurodegenerative disorders [13], they can also have protective functions. We and others have shown anti-inflammatory [14] and antinociceptive effects [15,16] of sEVs derived from antigen-presenting cells.

Opioids have been reported to alter cargo composition in sEVs and studied in the context of substance use disorder (SUD) and HIV infection [17,18]. A few studies have investigated the impact of morphine treatment on sEV cargo composition. Exposure of astrocytes to morphine resulted in a significant increase in the number of sEVs released compared to control cells [19]. sEVs from morphine-treated astrocytes also showed alterations in miRNA composition and 15 of the upregulated miRNAs were linked to the inhibition of morphine-mediated phagocytosis in recipient microglial cells [19]. They also reported that morphine-mediated dysregulation of miRNA cargo in astrocyte-derived sEVs led to pericyte migration and loss of pericyte coverage, and this effect was mediated by miR-23a in a mu-receptor-dependent manner [20].

In rodents, the administration of opioids by subcutaneous morphine pellet implantation, subcutaneous injections, osmotic minipumps, or infusions through intrathecal catheters for 3–12 days results in antinociception typically on the first day, followed by a loss of this effect and progressive hyperalgesia to thermal or mechanical stimuli over the course of several days [4,21,22]. Previous studies have shown that subcutaneous pellet implantation produced prolonged and robust OIH [23-25] and dependence as measured by significant increases in somatic signs of withdrawal [26,27]. Here we investigated chronic morphine-induced changes in miRNA composition in sEVs from the serum of mice after a morphine treatment paradigm that induces hyperalgesia. We tested whether administering these sEVs intrathecally contribute to analgesia in morphine-naïve recipient mice and prophylactically in a mouse model of inflammatory pain.

## MATERIALS AND METHODS

### Mice

All the studies were performed following the NIH guidelines and the protocols were approved by the Institutional Animal Care and Use Committee of Drexel University College of Medicine and the University of Pennsylvania. Eight-to ten-week-old male and female C57BL/6J mice were purchased from the Jackson Laboratory (Bar Harbor, ME). Mice were housed in a 12-hour light/dark cycle and provided with food and water *ad libitum*.

### Mouse models of pain and behavioral testing

For the OIH model by morphine pellet implantation, a morphine (75 mg morphine base) pellet or placebo (cellulose) pellet (NIDA Drug Supply Program) was implanted subcutaneously (*s.c*.) on the dorsal surface of mice under general isoflurane anesthesia. Serum for isolating sEVs was collected six days after implantation of morphine or placebo pellet. For the OIH model by morphine injections, morphine (#15464, Cayman Chemical, Ann Arbor, MI) was re-suspended in dimethyl sulfoxide (DMSO) and diluted with sterile phosphate-buffered saline (PBS) to a concentration of 10 mg/mL. Mice were subcutaneously injected twice daily with 20 mg/kg on days 1–3, and 40 mg/kg on day 4 as previously reported [28]. Mice were sacrificed after confirming the development of hyperalgesia and serum was collected for sEV isolation. For complete Freund’s adjuvant (CFA) model of inflammatory pain, mice were injected with 20 μL of 50% emulsified CFA (Sigma-Aldrich, St. Louis, MO) in PBS subcutaneously into the plantar hind paw.

Mice were habituated in behavioral testing rooms at least four days before baseline tests. The experimenters were blinded to the treatment conditions. Mechanical sensitivity was determined using the von Frey test. Static mechanical paw withdrawal thresholds were assessed by applying graded von Frey monofilaments to the plantar surface of the paw. The paw withdrawal threshold (PWT) was determined and evaluated by the up-down method [29]. Baseline measurements were taken for three days before any injection was administered and the values were averaged. The Hargreaves test was used to assess thermal sensitivity [16]. The baseline latencies were taken for three days before any injection was administered and were adjusted to approximately 10 seconds with a maximum of 20 seconds as the cutoff to prevent potential injury. Animals were placed on a plexiglass platform and a thermal beam was applied to the hind paw using an infrared light source under the plexiglass platform. The latency for the mice to withdraw their paws in pain was recorded as its threshold. Tail flick assays were performed as reported [30]. Briefly, the mouse tail was marked at three different points. The animal was restrained on the tail-flick test analgesia meter (Columbus Instruments, Columbus, OH), such that each marking is placed in the center of an intense light beam focused on the tail and the latency for the mice to flick its tail in pain was recorded as its threshold. The average of three different timings from each location on the tail was plotted. The latency at the first marking was done on all animals followed by the second marking on all animals then the third to ensure the accuracy of results. The heat intensity was adjusted to evoke a tail-flick baseline latency of ∼10 seconds. A cutoff time was set at 20 seconds to avoid tissue damage.

### Locomotor activity and behavior sensitization

Locomotor activity was analyzed in a home cage activity monitoring system (Med Associates Inc., St. Albans, VT). The testing cage, identical in dimension to the home cage, was placed in a photobeam frame (30 × 24 × 8 cm) with two levels of sensors arranged in an eight-beam array strip. For locomotor sensitization, animals were tested every 2–3 days for 120 min following an injection of saline or morphine (10 mg/kg, *s.c*.). On treatment days 1–3, all animals were administered saline. On days 4–9, animals received either morphine or saline, according to their treatment group. One μg sEVs were injected 24 hrs before Day 1. Beam break data was read into Med Associates personal computer-designed software and monitored at 10-min intervals.

### Conditioned place preference (CPP)

Morphine-naïve adult, male C57BL/6J mice (n = 8/group) underwent CPP training and testing to determine if sEVs derived from morphine-treated mice elicit a context-reward association as compared to sEVs derived from naïve mice. Training and testing took place in standard three-chamber CPP boxes with retractable doors (Med Associates Inc.). Each box consisted of one black chamber with grid floors, one white chamber with wire mesh floors, and a smaller central gray chamber with solid floor. The time spent and locomotor behavior in each chamber were automatically detected using photocell beam breaks, calculated by custom code for Med-PC V software. During a pre-conditioning session, mice were placed in the neutral chamber and allowed to freely explore all three chambers for 20 min. Using a biased design, the less-preferred chamber was assigned to be paired with sEV treatment. For the first conditioning session, all mice received an intrathecal injection of vehicle 1-hour prior to confinement in the vehicle-paired chamber for an hour. On the following day, mice were assigned based on initial preference to receive an intrathecal injection of vehicle, sEVs from naïve, or morphine-treated mice 1 hour prior to confinement to the sEV-paired chamber. This timing was selected to minimize the effects of the intrathecal injection itself on the development of a CPP and to reduce any carryover effect of sEV injection on subsequent conditioning sessions. Only a single sEV injection was used in order to reflect the behavioral paradigm. One day after the sEV-conditioning sessions, mice were returned to the boxes by an experimenter blinded to the treatment condition and were allowed to freely explore for 20 min in a test session identical to the pre-conditioning session. To control for initial side preference in this biased design, the CPP score was calculated by subtracting the time spent (sec) during pre-conditioning from the time spent during CPP assessment.

### Intrathecal sEV administration

Eight to ten-week-old C57BL/6J mice were used for intrathecal injections. All injections were performed with a Hamilton syringe and 30-gauge needles. After identification of the injection site, the needle was inserted into the tissue to the intervertebral space between L4 and L5, and a successful puncture resulted in a tail flick. Then 10–15 μL of 1 μg sEVs or sterile PBS was slowly injected.

### Ventral tegmental area (VTA) infusion of green PKH67 labeled sEVs

Fluorescent labeling of sEVs was done with general lipophilic fluorescent membrane dye, PKH67, according to manufacturer’s instructions (Sigma-Aldrich). Mice were anesthetized with isoflurane and placed in a stereotax. Mice received subcutaneous meloxicam (2 mg/kg) on the day of surgery. PKH67-labeled sEVs were injected in the mouse VTA (anteroposterior −3.4, lateral −0.3, dorsoventral −4.4;) [31].

### Isolation of sEVs from mouse serum

A combination of differential ultracentrifugation, ultrafiltration, and size-exclusion chromatography (SEC) was used to isolate sEVs from mouse serum. All centrifugations were performed at 4°C. Briefly, OIH model or control mice were anesthetized by isoflurane. Whole blood was collected via the eye in 1.7 mL microcentrifuge tubes without anticoagulant and incubated undisturbed at room temperature for 45 min to allow clot formation. The supernatant serum was collected after centrifugation at 2,000 × g for 10 min and stored at –80°C until use. The serum sample was diluted with an equal volume of Dulbecco’s phosphate-buffered saline (DPBS) without calcium and magnesium then centrifuged for 30 min at 2,000 × g to pellet cell debris. The supernatants were then centrifuged at 12,000 × g for 45 min and the suspension was filtered through a 0.22-μm syringe filter. Samples were then diluted to 4 mL with DPBS and transferred to 100 K Amicon Ultra Centrifugal Filters (Sigma-Aldrich), followed by centrifugation at 5,000 × g for 30 min. The concentrate was diluted to 500 μL with DPBS and purified by SEC using qEV original 35 nm Legacy columns (iZON, Medford, MA) following the manufacturer’s instructions. Four EV-rich fractions (7-10, 0.5 mL each) were pooled and after ultracentrifugation at 110,000 × g for 70 min (Optima TLX ultracentrifuge with TLA 100.4 rotor, Beckman Coulter, Inc.), the pellets were resuspended in DPBS and stored at –80°C until further use.

### Nanoparticle tracking analysis (NTA)

Brownian motion of sEV particles from mouse serum was visualized and size distribution and concentration of sEVs were measured using NanoSight NS300 (Malvern Instruments, United Kingdom). Briefly, sEV samples were diluted in DPBS and placed into sample chamber by syringe pump. The combination of shutter speed and gain were set to obtain 30-second videos which were analyzed by the NanoSight NTA 3.1.54 software.

### Detection of sEV surface epitopes by flow cytometry

The mouse MACSPlex exosome kit (#130-122-211, Miltenyi Biotec, Germany) was used to detect sEV specific markers using BD LSRFortessa (BD Biosciences, San Diego, CA) following the manufacturer’s protocol using 10 μg of sEVs as previously described [32]. Positive percentage (%Pos) of the IgG control was subtracted from each marker’s %Pos to remove background. To better represent both differences in percent positive and geometric mean fluorescent intensity (MFI), MFI for each marker was normalized to the highest observed MFI across the groups. Weighted expression was calculated as (%Pos < 60%) = % Pos, (Values >= %Pos 60%) = ((MFI/Highest MFI across groups) * % Pos * 0.5) + (Percent Positive * 0.5)), where 0.5 represents equal weighting of % Pos and weighted MFI expression. Samples are represented in the form of a heat map.

### miRNA profiling of sEVs and pathway analysis

Total RNA was isolated from sEVs using Ambion *mir*Vana miRNA isolation kit (Life technologies, Carlsbad, CA) following the manufacturer’s protocol. miRNA profiling was performed as described previously [33] using rodent Taqman low-density array (TLDA) microfluidic cards version A and B (Applied Biosystems, Foster City, CA) in 100 ng of total RNA. Undetermined cycle threshold (CT) values were first set to 40. Low expressed or undetected miRNAs with an average CT value of greater than 35 in all experimental groups were removed from the analysis. A miRNA was considered present in one experimental group only when the CT value of each replicate was lower than 35. Fold change was calculated from raw CT values using the 2 ^−ΔΔCT^ method [34]. The mean CT values of the top 10 miRNAs with the lowest standard deviations across all samples were used as the baseline CT for calculating the normalized ΔCT values. Statistical significance of differences in ΔCT values was determined by a two-tailed *t*-test for comparison of serum-derived sEVs from morphine-treated and control mice. A *p*-value threshold of 0.05 and an absolute fold-change of 2 were used to select significant differentially expressed miRNAs between experimental groups.

Genes targeted by differentially expressed miRNAs were identified using the union of the computationally predicted targets in TargetScan (with a Pct score of at least 0.90) [35] and the experimentally validated targets in miRTarBase [36]. Gene set enrichment analysis of the resulting targeted genes was done using ShinyGO V0.80 [37] to identify Gene Ontology (GO) biological processes and Kyoto Encyclopedia of Genes and Genomes pathways (KEGG). To reduce the redundancy among the reported GO biological processes, we applied a filtering where GO terms sharing more than 90% of their genes were removed, keeping only the GO terms with the most significant false discovery rate (FDR) value.

### Cell culture

HEK 293 and RAW 264.7 cells (American Type Culture Collection, ATCC, Manassas, VA) were grown in complete media (Dulbecco’s Modified Eagle Medium, DMEM, Corning, NY) supplemented with 10% heat-inactivated fetal bovine serum (FBS; Corning) and 1% penicillin/streptomycin (Gibco) at 37°C in 5% CO_2_.

### Luciferase reporter assay

The 3’untranslated region (UTR) of mouse *Creb1* (NM_133828.2) is 6912 base pair (bp) long, exceeding the capacity of the luciferase vector plasmid. Hence the *Creb* 3’UTR was cloned as 3 separate and overlapping fragments downstream of the luciferase gene to generate three constructs. HEK 293 cells were transfected with precursor miRNA for miR-155, miR-10a, or scrambled control, and luciferase reporter plasmid containing the 3’UTR plasmid using Lipofectamine 2000 (Life Technologies) for 48 hours. The Luc-Pair miR Luciferase assay kit (GeneCopoeia, Rockville, MD) was used to measure firefly and Renilla luciferase activity according to the manufacturer’s protocol. Firefly luciferase measurements normalized to Renilla were used as a transfection control. The data is the average of three independent experiments.

### Overexpression of miRNAs in RAW 264.7 cells

Lipofectamine RNAiMax transfection reagent (Life Technologies) was used for miRNA transfections and performed according to manufacturer’s protocol as described before [33]. After 24 hours, cell pellets were washed with 1× PBS and resuspended in either RNA lysis buffer (*mir*Vana kit) containing 0.5 U/μL RNAsin Plus (Promega; Madison, WI) for RNA isolation or 1X radioimmunoprecipitation assay (RIPA) buffer containing protease inhibitor cocktail (Thermo Scientific; Waltham, MA) for protein analysis.

### Cellular RNA isolation and qPCR

Total RNA was isolated using the *mir*Vana RNA isolation kit. RNA concentrations were determined using a NanoDrop ND1000 spectrophotometer (NanoDrop Technology Inc). The Maxima cDNA synthesis kit (Thermofisher Scientific) was used to generate cDNA and 2 μL cDNA was used for mRNA qPCR analysis. *Gapdh* (#4352339E, Applied Biosystems) was used as the normalizer. The primer probe used for *Creb1* was Mm00501607_m1 (Applied Biosystems).

### Western blotting

Cell or sEV samples were re-suspended in RIPA buffer (1:1) (Sigma-Aldrich) with protease inhibitors (1:100). Protein concentrations were measured by Micro BCA Protein Assay kit (Thermo Scientific) for sEVs or DC protein assay (Bio-Rad, CA, USA) for cell lysates. Equal amounts of protein samples in Laemmli SDS sample buffer (Thermo Scientific) were loaded on 10% Tris-Glycine gel (Thermo Scientific) and the gel was run at 125 V for 90 min. Proteins were transferred to 0.4-μm PVDF membrane at 25 V for 90 min, followed by blocking the membrane with 5% milk in Tris-buffered saline with 0.1 % Tween-20 (TBST) blocking buffer for 1 hour at room temperature. The membranes were incubated with primary antibodies in TBST and 10% (v/v) blocking buffer on shaker overnight at 4°C. The blots were washed with TBST thrice, 5 min each, then the blots were incubated with secondary antibodies in TBST at room temperature for 1 hour on the shaker. The membranes were washed with TBST thrice and protein detected by LI-COR Image Studio Software (LI-COR Biosciences, Lincoln, NE). Primary antibodies used were rabbit anti-albumin (1:2000, #16475-1-ap, Proteintech), rabbit anti-CREB (48H2) antibody (1:1000, #9197S, Cell Signaling), mouse anti-beta-actin (8H10D10) antibody (1:1000, #3700T, Cell Signaling). Secondary antibodies used were goat anti-rabbit HRP conjugate (1:10,000, #170-6515, Bio-Rad) and goat anti-mouse HRP conjugate (1:10,000, #170-6516, Bio-Rad). The blots were developed using SuperSignal west Femto substrate (Thermo Fischer).

### Statistical analysis

Data are presented as mean ± the standard error of the mean (SEM) from three or more independent experiments. Student *t*-test was used to determine the statistical significance for *in vitro* studies. Treatment effects were analyzed with a one-way analysis of variance (ANOVA) and the multiple comparisons between means were tested by the *post-hoc* Bonferroni method. All behavior data were analyzed using two-way ANOVA, and pairwise comparisons between means were tested by the *post-hoc* Šidák test. The differences between groups were considered significant when the *p*-value was less than 0.05. GraphPad Prism software (versions 10 and 11) was used for all statistical analysis.

## RESULTS

### Effect of chronic morphine treatment on mechanical and thermal threshold

To study the changes in sEV composition under morphine-induced hyperalgesia compared to control sEVs, we isolated sEVs from mouse serum. Since subcutaneous morphine pellet implantation and repeated morphine injections are both used to generate OIH models, we investigated the development of chronic morphine-induced hyperalgesia using both methods. Subcutaneous morphine pellet implantation causes prolonged and robust OIH [23-25]. The study schematic following 75 mg morphine, or a placebo (cellulose) pellet implantation is shown in Fig. 1A. As expected, morphine induced antinociception with an increase in mechanical threshold in the von Frey test, but this effect was lost by day 4 (Fig. 1B). Assessment of thermal sensitivity as determined by the Hargraves test (Fig. 1C) and tail flick assay (Fig. 1D) showed highest paw withdrawal latency times on day one, which diminished gradually. This was followed by a decrease in paw withdrawal latency comparable to the control group. In mice that received subcutaneous morphine injections for four days, [28] (Fig. 1E) mechanical allodynia (Fig. 1F) and thermal hyperalgesia (Fig. 1G) developed on day 5, and thermal hyperalgesia lasted for 12 days. Overall, our studies confirmed previous observations [25] that continuous morphine treatment induced mechanical allodynia and thermal hyperalgesia.

**Fig. 1.**
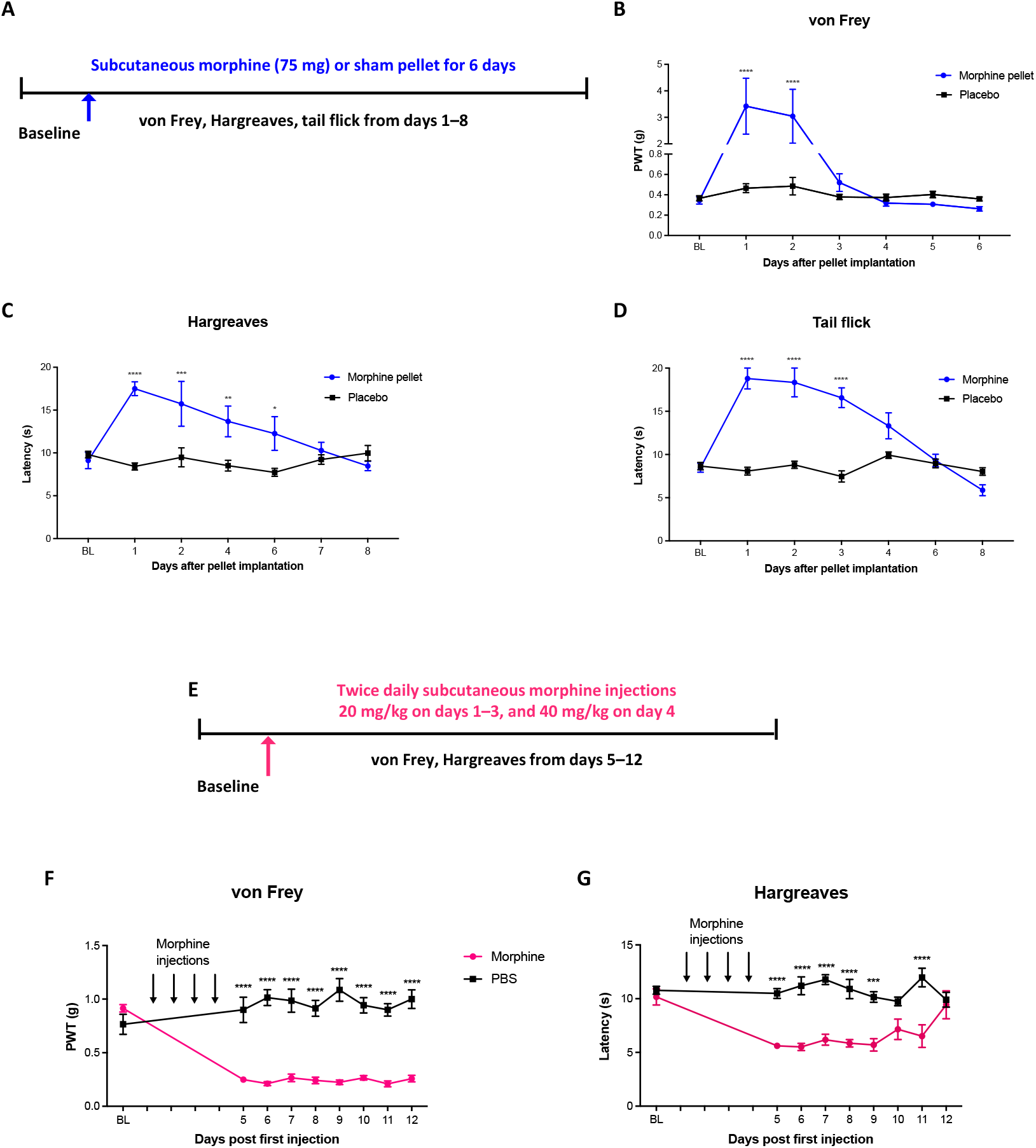
Morphine induced OIH in mice. **A** Schematic representation of chronic morphine-induced hyperalgesia model using subcutaneous implantation of 75 mg morphine or a placebo pellet. **B** Mechanical allodynia determined by von Frey test, thermal hyperalgesia assessed using **C** Hargreaves test and **D** tail flick assay in mice implanted with morphine pellet. **E** Schematic of escalating doses of morphine (20 mg/kg twice daily for 3 days and 40 mg/kg on the 4th day, *s.c*.) model for OIH. **F** Mechanical allodynia and **G** thermal hyperalgesia in mice receiving escalating doses of morphine. Ten-week-old male C57BL/6J mice were used (n=7). Statistical analysis was determined by two-way repeated measures ANOVA followed by Šidák’s multiple comparisons test; \**p*<0.05, \*\**p*<0.01, \*\*\**p*<0.001, \*\*\*\**p*<0.0001. PWT, paw withdrawal threshold. BL, baseline.

### Characterization of serum-derived sEVs

sEVs were characterized for the presence or enrichment of endosome-derived membrane proteins such as tetraspanins and assessed using bead-based flow cytometry. In addition to confirming the expression of tetraspanin proteins, we also observed that though the sEVs from the serum of morphine-treated (Mor-sEVs) and naïve mice (Naïve-sEVs) had a similar expression, the expression of CD146, CD69, and CD105 were lower in Mor-sEVs (Fig. 2A). We also confirmed the purity of sEVs by western blot analysis for the negative control protein albumin which is detected in the serum but absent in sEVs (Fig. 2B). We performed nanoparticle tracking analysis (NTA) of serum-derived sEVs and observed the mean concentrations of naïve-sEVs and mor-sEVs to be 12.36 ± 2.01 × 10^8^ and 10.25 ± 0.85 × 10^8^ particles/mL, respectively (Fig. 2C). The mean diameters were less than 150 nm (Fig. 2D), indicating these vesicle preparations fall within the size range of sEVs. There was no statistical difference between sEVs from controls and sEVs from morphine-treated mice.

**Fig. 2.**
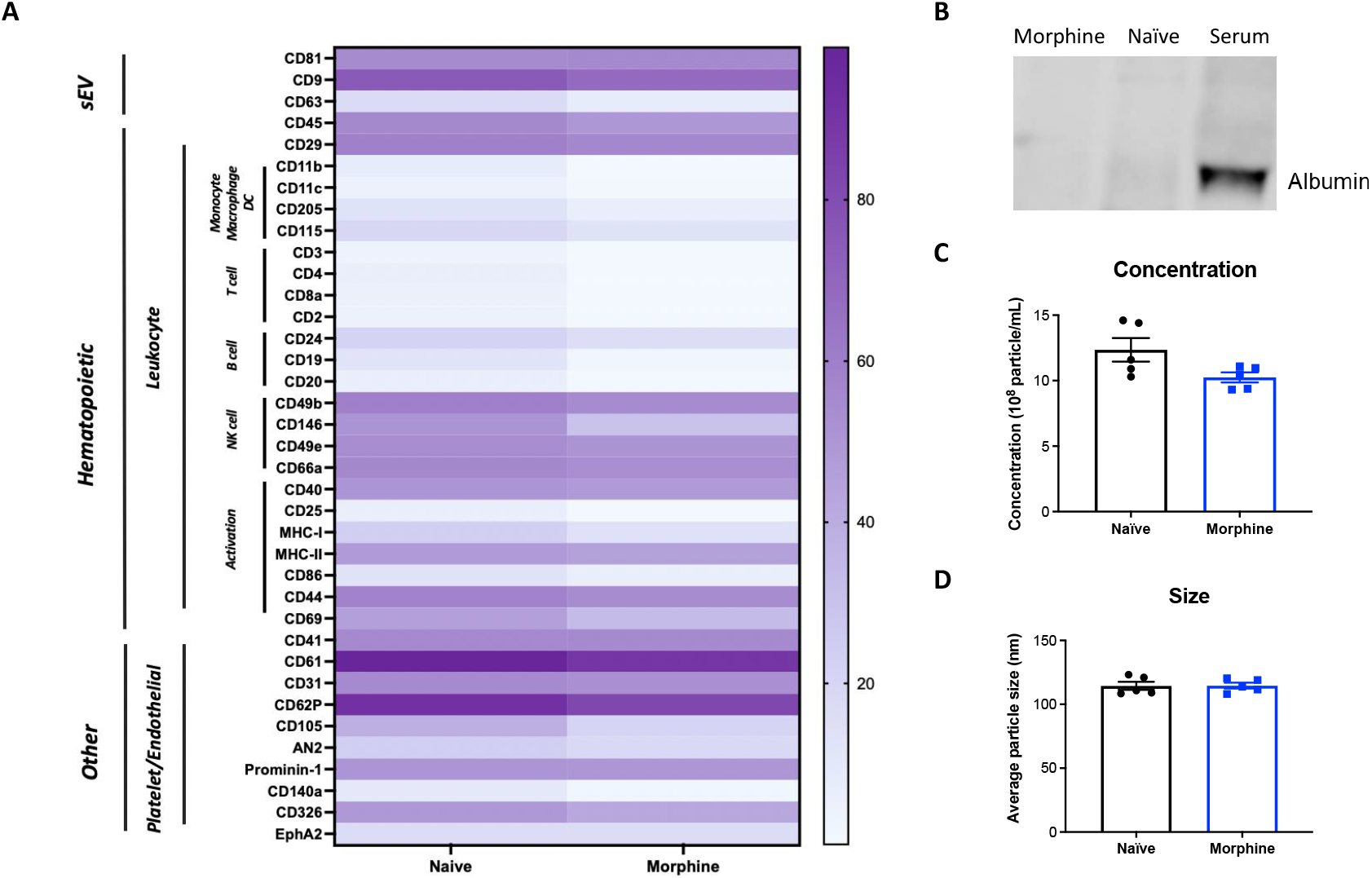
Characterization of sEVs purified from mouse serum. **A** Characterization of markers on serum-derived sEVs from naïve and OIH mice using a MACSPlex flow cytometry bead-based capture and fluorescent detection assay. sEVs (n=1) used were from a pool of serum of 5 mice of each treatment. Comparisons are normalized and weighted to represent alterations in both percent positive and fluorescent intensity. **B** Western blot analysis of sEV lysate indicates that albumin is absent in sEV samples, indicating purity. **C** Nanoparticle tracking analysis (NTA) of naïve- and morphine-derived sEVs indicated mean concentrations of 12.36 ± 2.01 × 10^8^ and 10.25 ± 0.85 × 10^8^ particles/mL, respectively; and **D** mean diameters of 114.48 ± 7.05 and 114.58 ± 5.15 nm, respectively, indicating purity.

### miRNA composition differs in sEVs purified from serum of morphine-treated mice vs control mice

We investigated chronic morphine-induced changes in miRNA composition of sEVs in the serum. Both male (n=5/group) and female (n=4/group) mice were included to identify sex-specific differences. Profiling of 758 miRNAs by qPCR (Supplementary Table S1) showed significant differential expression of 18 miRNAs in sEVs from morphine treated mice compared to control. The Venn diagram summarizes the number of sex-specific miRNAs that are significantly altered (Fig. 3A) and the number of miRNAs up or downregulated in Mor-sEVs are shown (Fig. 3B). Presence-absence analysis showed that all the miRNAs expressed in the female OIH model were found in the female control (Fig. 3C, Supplementary Table S2). However, the male OIH model had 15 exclusively present and 16 exclusively absent miRNAs compared to the same-sex control. These suggest that the male OIH model has a specific miRNA signature. There are 37 commonly expressed miRNAs across all the groups. Fig. 3D shows miRNAs upregulated in Mor-sEVs and Supplementary Table S3 lists all significantly altered miRNAs and the corresponding fold changes and p values.

**Fig. 3.**
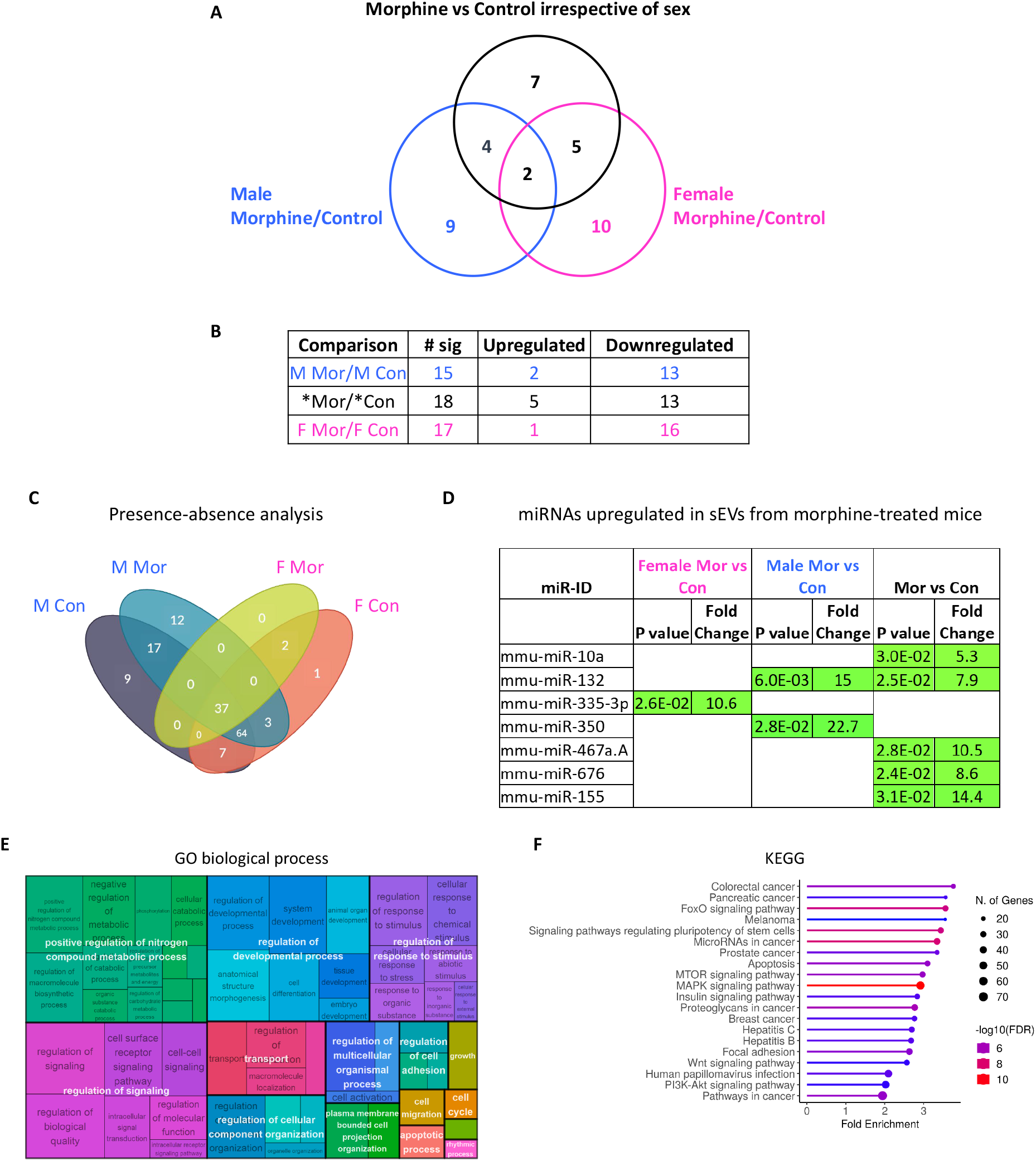
Differentially expressed sEV miRNAs in morphine pellet treated mice. **A** Venn diagram showing relative expression of miRNAs from serum sEVs of 4 male (M) and 4 female (F) Mor-treated mice compared to control. Comparison of miRNAs shown between all morphine pellet treated mice vs all controls irrespective of sex (*mor/*con) and between sex-specific groups. Significance was determined using a two-tailed *t*-test for comparison of morphine- or control-derived samples. A *p*-value threshold of 0.05 and an absolute fold-change of 2 was used to identify differentially expressed miRNAs between groups. **B** Summary of sex-specific comparison of sEV miRNAs listing the number of significantly different miRNAs and directionality of changes. **C** The presence and absence analysis of miRNAs in sEVs from male and female morphine treated and control mice. Morphine treatment-induced, sex-specific, and commonly-expressed miRNAs across groups shown. **D** miRNAs upregulated in sEVs from morphine-treated mice. The *p* value and fold change of miRNAs including miR-10a and miR-155 predicted to bind *Creb1* mRNA is shown. The significance was determined using two-tailed *t*-test for comparison of morphine or sham pellet implanted samples. A *p*-value threshold of 0.05 and a fold-change of 2 were used to select significantly differentially expressed miRNAs between experimental groups. **E** Gene Ontology (GO) biological processes and **F** KEGG pathways that are significantly enriched (FDR < 0.01) with the genes that are targeted by the differentially expressed miRNAs. Data were analyzed from mice of both sexes with n=4/group.

Biological processes and pathways regulated by the differentially expressed miRNAs irrespective of sex were identified by compiling the target genes of these miRNAs followed by gene set enrichment analysis. Major categories of enriched biological processes included regulation of nitrogen metabolism, developmental processes, response to stimulus and signaling (Fig. 3E, Supplementary Table S4). Pathway enrichment analysis indicated that the genes regulated by the miRNAs that are differentially expressed in morphine-treated mice are also involved in various cancers and signaling pathways including mTOR, MAPK, Wnt, and PI3K-Akt (Fig. 3F). Supplementary Table S5 and Supplementary Fig. 1 GO show the subsets of the significantly enriched biological processes that CREB1 is involved in.

### Confirmation of miRNA binding to the 3′ untranslated region (3’UTR) of *Creb1* and transcriptional regulation of target genes

Small noncoding miRNAs regulate gene expression by binding predominantly to the 3′UTR of mRNAs by 6-to 8-bp seed sequence complementarity. We performed bioinformatic analysis of miRNAs upregulated in sEVs from morphine treated mice and observed that several of the miRNAs are predicted to target *Creb1* mRNA. Since there were multiple miRNAs predicted to target *Creb1* 3’UTR, including some miRNA target sites represented more than once, we selected two miRNAs, miR-155 and miR-10a, that were significantly upregulated in the morphine group as a whole (male and female mice combined) versus the control group (Fig. 3D). miR-132 is another miRNAs that we identified and previous studies have validated miR-132 to bind and negatively regulate *Creb1* [38].

Upon binding, miRNAs can induce either mRNA degradation or translational repression and thus negatively regulate the expression of target genes [39,40]. To confirm miRNA binding to the 3’UTR of *Creb1*, we performed a luciferase reporter assay. Since the *Creb1* 3’UTR is ∼7 kb, we cloned it as 3 separate fragments (*Creb1-1, −2* and *-3*, schematic shown in Fig. 4A) to test them for miRNA binding in the reporter assay. Fig. 4B shows the reduction in luciferase activity 48 hours after transfection of miR-10a and miR-155. miR-10a has binding sites present in fragment ‘2’ and ‘3’ but more efficient binding was observed in the fragment ‘3’ compared to the fragment ‘2’.

**Fig. 4.**
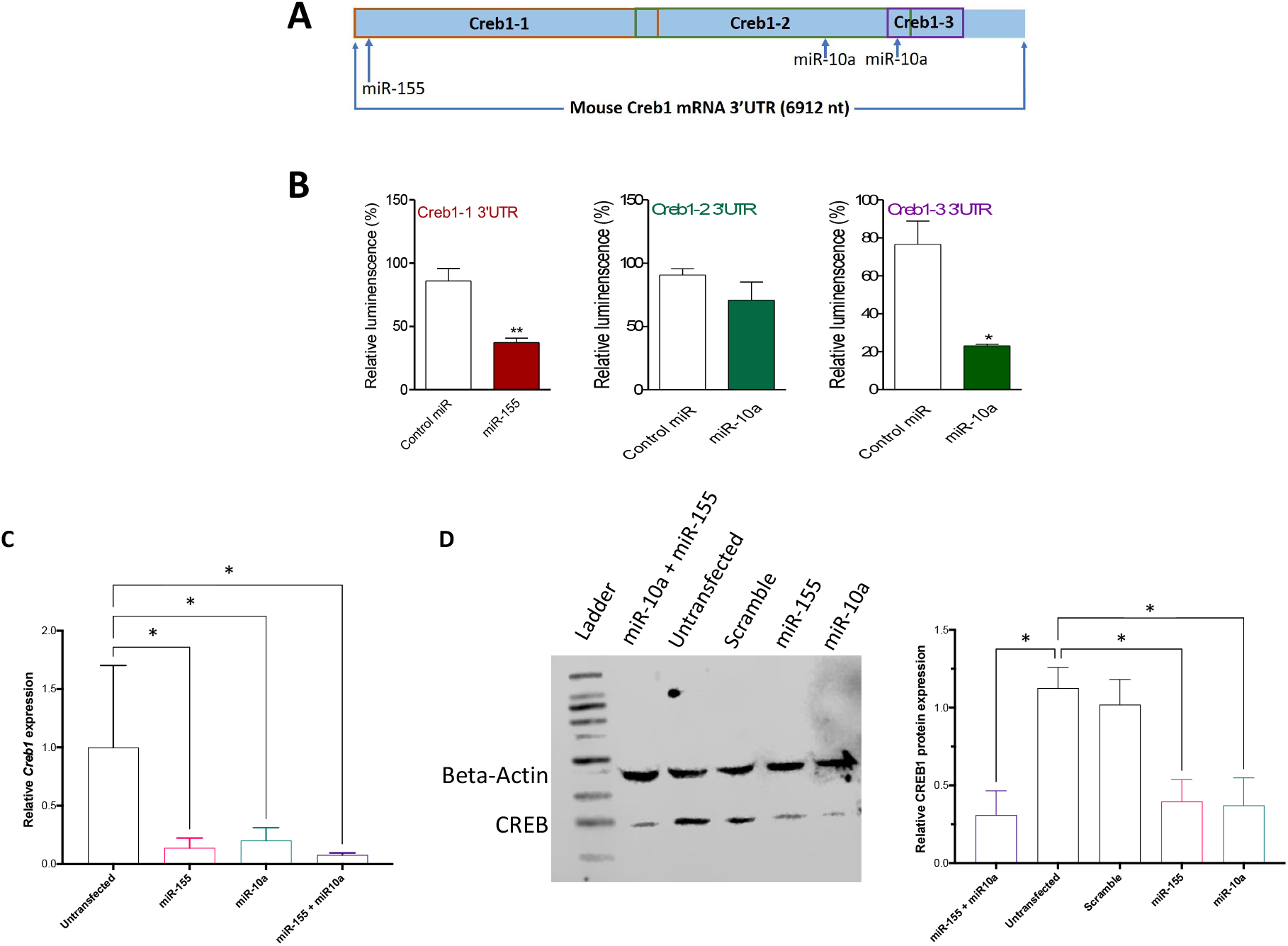
Regulation of *Creb1* expression by miR-155 and miR-10a. **A** Schematic representation of miRNA target sites on *Creb1* mRNA 3’UTR and the regions corresponding to 3’UTR used in luciferase assay to validate mRNA-miRNA binding. **B** Confirmation of miR-155 and miR-10a binding to the 3’UTR of *Creb1*. Reporter assay shows a significant reduction in relative luminescence in cells transfected with miR-10a or miR-155 in comparison to control miRNA, indicating that miR-155 and miR-10a can bind to *Creb1* 3’UTR. \**p*<0.05, \*\**p*<0.005, two-tailed unpaired *t*-test. **C** Taqman analysis of endogenous *Creb1* in RAW 264.7 cells 24 hours after transfection of miR-155, miR-10a or both miRNAs. **D** Western blot analysis and quantification 24 hours after transfection of RAW 264.7 cells with miR-155, miR-10a or both miRNAs, significantly reduced CREB1 protein. Significance was determined using Student’s *t*-test, *p*-value *<0.05, **<0.01, ***<0.001.

To test the ability of miR-155 and miR-10a to modulate endogenous *Creb1* mRNAs *in vitro*, we transfected mouse RAW 264.7 macrophage cells with miRNA mimics or scrambled control miRNA. Since miRNAs can regulate gene expression by mRNA degradation or translational repression, we performed qPCR to measure endogenous levels of target mRNAs, 24 hours after transfection. There was a reduction in *Creb1* mRNA levels upon miRNA transfection suggesting that miR-155 and miR-10a binding can downregulate *Creb1* (Fig. 4C). To further confirm that miR-155 and miR-10a can negatively regulate protein expression of CREB1, a western blot analysis of RAW 264.7 macrophage cell lysate 24 hours after miRNA transfection was performed. Fig. 4D shows that overexpression of these miRNA mimics resulted in lower levels of CREB protein and there was an additive effect upon transfection with both miRNAs which was more evident in the qPCR (Fig. 4C).

### Prophylactic intrathecal administration of sEVs isolated from chronic morphine-treated mice attenuates thermal hypersensitivity

To investigate the role of sEVs released during chronic morphine exposure in modulating pain, we tested if exogenously administering 1 μg of sEVs from morphine treated or control mice, intrathecally into naïve male recipient mice can modulate basal pain thresholds. A schematic of paradigm is shown in Fig. 5A. We tested the effect of sEVs on basal threshold using sEVs from both morphine pellet implanted (Fig. 5 B-E) and morphine injected (Fig. 5 F-J) models along with their respective control mice. We observed that 1 μg of mouse serum-derived sEVs obtained from morphine pellet implanted-male sEV OIH model donor mice when administered intrathecally into drug-naïve male recipient mice resulted in an increase in thermal (Fig. 5C), but not mechanical (Fig. 5B), paw withdrawal latency. This indicates elevated pain thresholds on day 3 in mice treated with Mor-sEVs from pellet model OIH mice. However, Mor-sEVs from morphine injection model mice did not alter basal pain thresholds in recipient mice (Fig 5F and G). We also tested the efficacy of sEVs from morphine injection OIH model mice in a tail flick model to assess the analgesic effect to thermal stimuli and did not observe any significant difference at 3 hours post sEV injection (Supplementary Fig. 2). We then investigated how prophylactic administration of these sEVs impacted nociceptive threshold to a future painful stimulus. Inflammatory pain was induced by an intraplantar injection of CFA 10 days after sEV administration. Irrespective of the method (morphine pellet or injection) used to induce OIH, mice injected with Mor-sEVs showed attenuation of CFA-induced thermal hypersensitivity (Fig. 5E and I). There was also a significant attenuation of CFA-induced mechanical hypersensitivity in mice treated with Mor-sEVs from the OIH injection model as compared to control (Fig. 5H), but not in the mice that received OIH pellet model-derived sEVs (Fig. 5D).

**Fig. 5.**
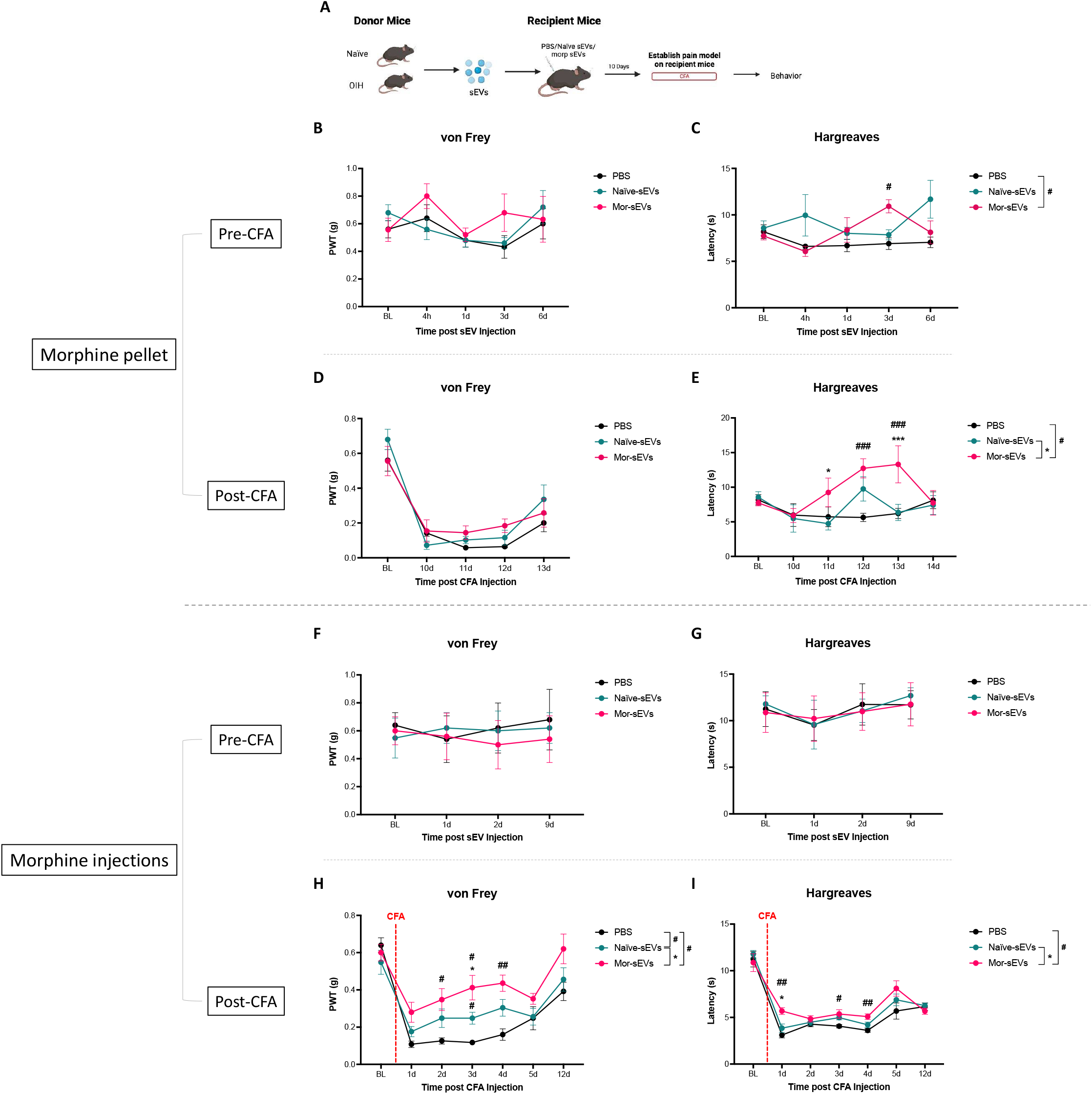
Prophylactic intrathecal administration of sEVs isolated from the serum of chronic morphine-treated mice attenuates pain hypersensitivity. **A** Schematic representation of experimental design. After baseline behavioral testing, 1 μg sEVs from male naïve or morphine-treated donor mice or PBS were injected intrathecally into nine-week-old male recipient mice. Mor-sEVs from morphine pellet model OIH mice did not alter basal **B** mechanical thresholds but increased basal **C** thermal thresholds 3 days after administration. CFA model was established in recipient mice 10 days post sEV injection. Mechanical and thermal thresholds were assessed in recipient mice with sEVs from **D-E** morphine pellet and **F-G** repeated morphine injection OIH model mice. Statistical analysis was performed by repeated measures two-way ANOVA, data shown are mean ± SEM (n=5) *\*, #p*<0.05; *\*\**, ##*p*<0.01; ***, ###*p*<0.001. Mor, morphine. PWT, paw withdrawal threshold. BL, baseline.

### Behavior response to morphine administration in sEV-treated mice

Behavioral sensitization serves as an important model of neural plasticity and has been suggested as a mechanism that may model aspects of human drug addiction [41]. We have previously used locomotor activity to study the sensitizing effects of morphine [42-44]. Since acute morphine administration can elevate locomotor activity [43], we investigated if administration of sEVs from morphine treated or control mice can impact locomotor activity in naïve recipient mice. We first confirmed successful injection of sEVs directly into the ventral tegmental area (VTA). For this, sEVs were labeled with PKH67 green fluorescent membrane dye and Fig. 6A-C show successful infusion of sEVs. Our results show that injection of Mor-sEVs from OIH pellet model do not impair the normal development of behavioral sensitization to morphine (Fig. 6D) in naïve sEV recipient mice.

**Fig. 6.**
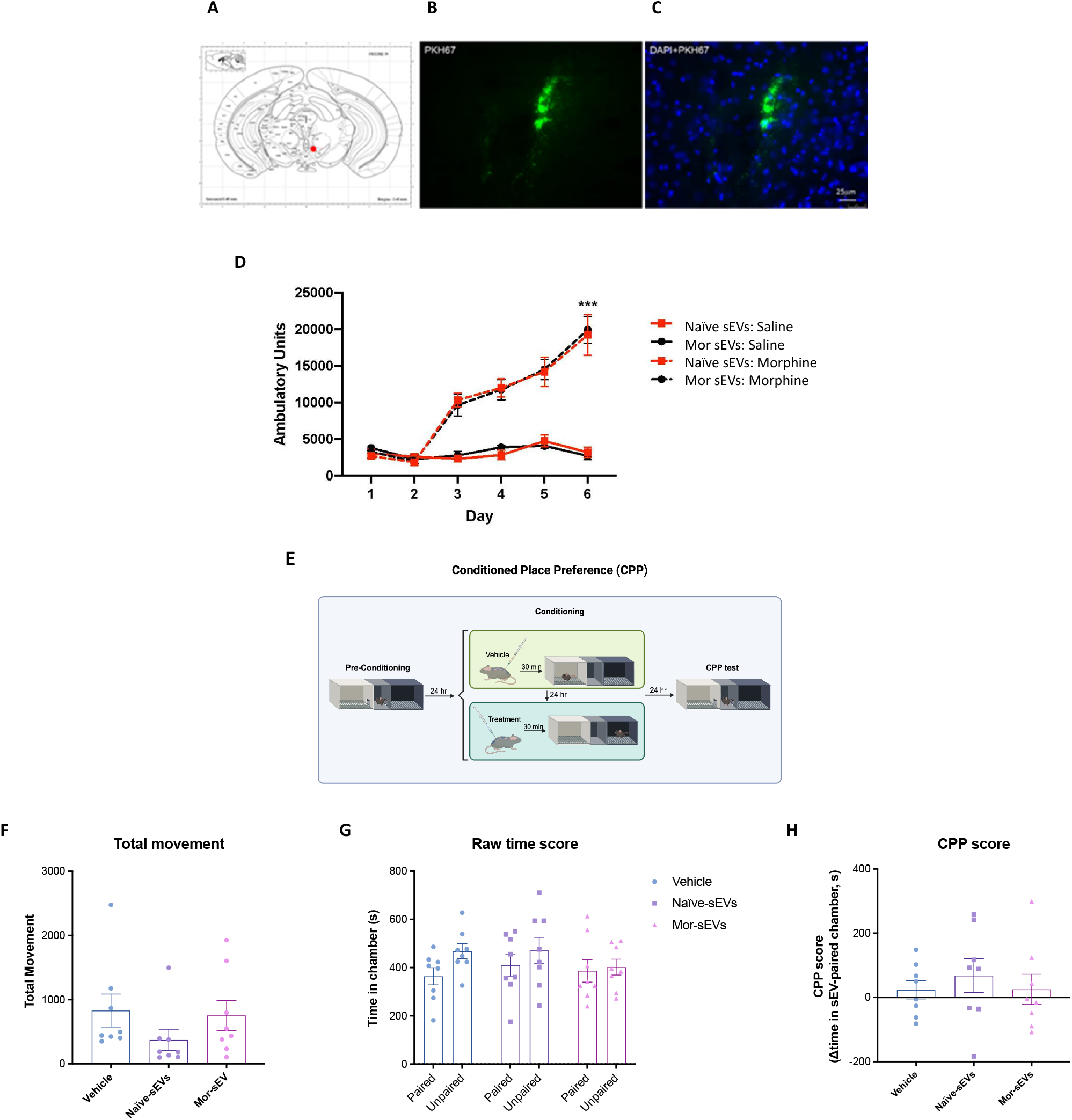
Treatment with sEVs does not impair morphine-induced locomotor sensitization and does not induce a conditioned place preference. Detection of PKH67-labeled sEVs in the mouse VTA (anteroposterior −3.4, lateral −0.3, dorsoventral −4.4) 1 hour after infusion. **A** Schematic representation of injection site. **B** PKH67 green label **C** PKH67 co-stained with DAPI (blue). Images taken at 40X magnification. **D** sEVs from morphine pellet implanted OIH model injected directly into the ventral tegmental area do not impair morphine-induced locomotor sensitization in C57BL/6J male mice. Saline was administered to both groups on days 1–3 and morphine (20 mg/kg) on days 3–6. The saline group received saline on all days, n=6/group. *\*\*\*p<*0.001, main effect of group and of morphine day 3 compared to day 6. **E** Schematic representation of conditioned place preference paradigm. **F** Total movement did not significantly differ between groups, suggesting sEVs from morphine injection model of OIH do not impair movement. **G** Total time in chambers and **H** CPP scores did not differ between vehicle or sEV-treated mice. No groups exhibited a CPP score significantly different from zero, suggesting that sEV-treatment did not elicit a CPP. Statistical analysis was performed by one-way ANOVA, bars represent mean +/- SEM (n=8).

### sEVs do not induce a conditioned place preference

We have previously reported the successful of delivery of dye-labeled sEVs to the spinal cord and DRG after intrathecal injections [16,45]. To determine whether sEVs from morphine OIH model impacted locomotor behavior, the movement during the CPP and conditioning sessions was measured by consecutive beam breaks. The CPP paradigm is described in Fig. 6E. A repeated measures ANOVA (treatment day vs. treatment group) indicated a main effect of treatment day, but no effect of treatment group or group x day interaction, suggesting that sEVs did not significantly impact movement (Fig. 6F). Since overall movement was reduced on the second day of injections regardless of treatment group, this suggests that it was likely an effect of consecutive intrathecal injections (Supplementary Fig. 3). To determine whether sEV injection induced a preference for sEV-paired chambers, total time spent in each chamber was compared (Fig. 6G). As a biased design was used, CPP scores were calculated as time spent in the sEV-paired chamber after conditioning minus time spent prior to conditioning. No significant effect of sEV treatment was observed on CPP scores (one-way ANOVA) (Fig. 6H). Further, CPP scores did not significantly differ from zero for any of the treatment groups (one sample t-test) indicating that mice did not form a preference for sEV-paired chambers.

## DISCUSSION

All cells release sEVs and these vesicles enable a cohort of biomolecules, including miRNAs, to travel long distances via circulation. Upon uptake, sEVs can modulate gene expression in recipient cells. Previous reports have shown that chronic morphine exposure can induce prolonged and robust OIH [23-25] as well as opioid dependence as measured by significant increases in somatic signs of withdrawal [26,27]. Thus, we chose a chronic morphine treatment paradigm to enable studies on the role of OIH model-derived sEVs. We hypothesized that there would be distinct sEV miRNA signatures in prolonged morphine-treated mice which may be involved in mediating the signaling related to SUD and pain. Differential expression of miRNAs in males and females has been reported [46-48]. In combining miRNAs detected in male and female mice, we found significant differential expression of 18 sEV miRNAs in morphine-treated group compared to control mice.

Pathway enrichment analysis for miRNAs differentially expressed in morphine-treated mice implicated several pathways including mTOR, MAPK, Wnt, and PI3K-Akt signaling with MAPKs or mitogen-activated protein kinases having the largest representation. MAPKs have a critical role in the signal transduction cascade associated with peripheral and central sensitization linked to the initiation and maintenance of chronic pain [49]. Activation of the MAPK signaling pathway leads to the phosphorylation and activation of CREB. This, in turn, results in the transcription of pain-related genes that contribute to the development and maintenance of chronic pain states. Increased CREB activation in neurons has been associated with heightened pain perception and the development of chronic pain. For instance, in models of chronic pain, CREB phosphorylation in the dorsal horn neurons, and peripheral sensory neurons correlates with increased pain sensitivity. Phosphorylation of MAPKs and CREB is involved in the morphine-induced increase in spinal pain-related neuropeptides, calcitonin gene-related peptide (CGRP) and substance P, levels in primary sensory afferents, contributing to the development of tolerance to opioid-induced analgesia [50]. Also, chronic opioid use activates mTOR signaling, contributing to the development of hyperalgesia; repeated intrathecal administrations of morphine resulted in spinal dorsal horn mTOR activation through μ opioid receptor-triggered PI3K/Akt signaling [51]. Wnt proteins, through both canonical (β-catenin-dependent) and non-canonical pathways, contribute to pain sensitization, neuronal excitability, and neuroinflammation [52]. β-catenin can indirectly activate CREB by regulating signaling cascades such as PI3K/Akt and MAPK, both of which lead to CREB phosphorylation. Thus, Wnt signaling could enhance CREB-mediated transcription of genes involved in pain such as *BDNF*, contributing to chronic pain states. Wnt signaling also plays a critical role in withdrawal symptoms from opioid receptor activation in mice [53]. miRNAs altered in blood from multiple rodent models of pain, including neuropathic, inflammatory, and chemotherapy-induced pain models are predicted to target *Wnt* genes [54] suggesting downregulation of miRNAs that target Wnt may be a common mechanism in regulating pain.

Since miRNAs are negative regulators of gene expression, we performed bioinformatic analysis [55] focusing on miRNAs upregulated in sEVs from morphine-treated mice. Several of the miRNAs significantly altered in sEVs from morphine-treated mice that we identified were bioinformatically predicted to target *Creb1* 3’UTR. We observed that Creb1 is a validated target of miR-132 [38,56,57]. We chose to focus on two miRNAs, miR-10 and miR-155, that were significantly upregulated in the morphine group compared to the control group. CREB is a well-characterized transcription factor implicated in underlying mechanisms of both pain and drug addiction-related behaviors. A variety of addictive drugs have been shown to alter the expression and/or activation of CREB and alterations in CREB activity are linked to behavioral manifestations of opioid reward, tolerance and withdrawal [26,27,42,58]. Morphine can alter the levels and function of CREB, and CREB signaling mediates a variety of morphine-induced behaviors. However, the mechanism of morphine induced CREB regulation has been less well studied. We confirmed miRNA binding to the *Creb* 3’UTR by luciferase reporter assay. Binding sites for miR-10a are present in fragments ‘2’ and ‘3’ that we generated for the ∼7kb long *Creb* 3’UTR, but we observed more efficient binding of miRs to fragment ‘3’ compared to fragment ‘2’. Target locations are not evenly distributed throughout the whole UTR and there is a strong preference for targets located in close vicinity of the stop codon and the polyadenylation sites [39]. This could contribute to the differences observed for miR-10a in our luciferase assays. Additionally, the role of RNA secondary structure in blocking miRNA access also cannot be ruled out. Down-regulation of endogenously expressed mRNA and protein following miRNA transfection indicates that changes in CREB expression can be mediated by mRNA degradation or translational repression.

Studies have shown significant differences between the populations of miRNAs in cells and their secreted sEVs; certain miRNAs are selectively sorted into sEVs thereby lowering their expression levels in the cells secreting them, while other miRNAs are selectively retained by cells [59-61]. Thus, miRNA availability for secretion in sEVs is controlled, at least in part, by the cellular levels of their targeted transcripts [62,63]. Our previous studies using cultured postnatal mouse cortical neurons and astrocytes showed a distinct miRNA profile between sEVs and their corresponding source cells. We observed that only 20.7% of astrocyte-derived miRNAs were loaded into sEVs, while 41.0% of neuron-derived miRNAs were loaded into sEVs [64], indicating differences and specificity in the cellular sorting mechanisms and that cargo loading into sEVs is a highly regulated process. Our results here demonstrate that chronic morphine alters sEV composition. Previous miRNA profiling of sEVs from morphine-treated astrocytes showed enrichment of miRNAs with an AU- or GU-rich motif [19]. Since it is challenging to determine the source of sEVs in circulation, future studies will determine how morphine treatment can impact this packaging along with the role of specific motifs and RNA binding proteins in determining selective sorting of miRNAs into sEVs from cultured central nervous system cells.

To investigate the functional impact of transferring sEVs from OIH model to naïve mice, we intrathecally injected 1 μg of Mor-sEVs into naïve recipient mice. Mor-sEVs from morphine pellet implanted but not morphine injection model of OIH donor mice resulted in an elevated thermal pain threshold in naïve recipient mice on day 3. sEVs from control mice did not induce any alterations in basal mechanical or thermal thresholds at 1 μg as we reported previously [32]. We next assessed if prophylactic Mor-sEVs can induce faster resolution of inflammatory pain as observed previously with 1 μg RAW 264.7 macrophage cell line derived sEVs [16] and 10 μg of mouse serum derived sEVs (naïve and neuropathic pain model mice) [32]. Interestingly 1 μg of serum-derived Mor-sEVs from both morphine pellet and injection models of OIH showed an earlier resolution CFA-induced thermal hypersensitivity in naïve recipient mice. Mor-sEVs from morphine injection OIH model also resulted in earlier resolution of mechanical hypersensitivity. Collectively, our studies show that 1 μg sEVs from OIH model was more efficacious in inducing behavioral changes in recipient mice compared to sEVs from the serum of naïve control mice, or spared nerve injury model of neuropathic pain [32]. How different morphine treatment paradigms impact the composition of miRNAs and other sEV cargo in circulation and how the uptake of these sEVs by recipient cells contribute to behavioral responses underlying inflammatory pain remain to be elucidated. Pathway analysis of miRNAs altered in blood from rodent models of pain, [54] suggest there could be shared mechanisms such as regulation of Wnt signaling.

We used two well-established behaviors to determine if administering sEVs from morphine-treated mice alters addiction-like behaviors including locomotor sensitization and CPP in recipient mice. We found that administering sEVs from morphine-treated mice does not impair the normal development of behavioral sensitization in recipient mice and sEVs do not impact subsequent response to morphine in recipient mice. We used sEVs from morphine pellet implanted OIH model for locomotor sensitization test and since we did not see an effect, we tested sEVs from the repeated morphine injection model for the CPP test. These sEVs did not induce a CPP in recipient mice. We also did not observe any aversive learning associated with sEV administration suggesting that sEVs did not induce conditioned place aversion.

We used male and female mice for miRNA profiling, but all the recipient mice used were male. Morphine is more potent in males compared to females and this sex differences in opioid analgesia have been attributed to higher levels of mu opioid receptor expression and binding in males [65]. Future studies matching sexes of sEV donor and recipient mice will help elucidate if there are sex specific differences in sEV function. Another limitation to our studies is that we used only 1 μg sEVs and this potentially resulted in modest behavior effects in recipient mice. Using higher concentrations of sEVs may produce more robust behavior outcomes in sEV recipient mice. A major challenge for successful translation remains the difficulty to precisely target the cell type or organ of interest whilst limiting off target biodistribution. We have previously reported the presence of endogenous opioids, specifically leu-enkephalin in serum derived sEVs [32]. It will be interesting to investigate how morphine treatment impacts the packaging and thus the sEV composition of endogenous opioids in OIH models. In sum, our data indicate that administration of 1 μg sEVs from OIH models can induce desired behavioral outcomes *in vivo, i.e*., a protective role for sEVs from morphine-treated mice in attenuating pain hypersensitivity without augmenting behaviors associated with opioid use disorders. As CREB is not a “druggable” target, the potential to modulate its expression *in vivo* with sEVs could prove to be an innovative approach to regulate gene expression.

## ACKNOWLEDGEMENTS

We thank Julia K. Brynildsen for stereotaxic injections of sEV into the VTA and locomotor behavior assay

## FIGURE LEGENDS

**Supplementary Fig. 1** Map of the subset of Gene Ontology (GO) biological processes that CREB1 is involved in that are significantly enriched with the genes targeted by the differentially expressed miRNAs.

**Supplementary Fig. 2 sEVs from injection-model OIH mice do not alter basal thermal thresholds in the tail-flick assay**. Basal thermal thresholds via the tail flick assay, were not altered after Naïve- or Mor-sEV injections derived from OIH injection model donor mice. Statistical analysis was performed by repeated measures two-way ANOVA, data shown are mean ± SEM (n=5).

**Supplementary Fig. 3 sEVs do not alter total movement in CPP paradigm**. Total movement was reduced following a second intrathecal injection on the treatment day, but this was not mediated by sEVs. Statistical analysis was performed by one-way ANOVA, \*\*\**p*<0.001, bars represent mean +/- SEM (n=8).

**Supplementary Table S1** Raw and delta cycle threshold (CT) of miRNAs detected using Taqman low-density array microfluidic cards.

**Supplementary Table S2** List of miRNAs present in serum sEVs from mice of both sexes implanted with morphine or control pellets.

**Supplementary Table S3** List of differentially expressed miRNAs in male, female and combined group of morphine pellet implanted mice vs their respective control groups. Significance was determined using a two-tailed *t*-test for comparison of morphine or control derived samples. A *p*-value threshold of 0.05 and an absolute fold-change of 2 was used to identify differentially expressed miRNAs between groups. Orange-red rows represent downregulation, and green rows show upregulation of miRNAs.

**Supplementary table S4**. List of Gene Ontology (GO) Terms from biological process and molecular function that are significantly enriched with the genes that are targeted by the differentially expressed miRNAs in sEVs from morphine pellet treated mice compared to control.

**Supplementary table S5**. List of the subset of GO terms from biological processes that CREB1 is involved in that are significantly enriched with genes targeted by the differentially expressed miRNAs sEVs from morphine pellet treated mice compared to control.

