## Supplementary Fig for "Morphine-induced hyperalgesia impacts small extracellular vesicle miRNA composition and function"

### Slide 1
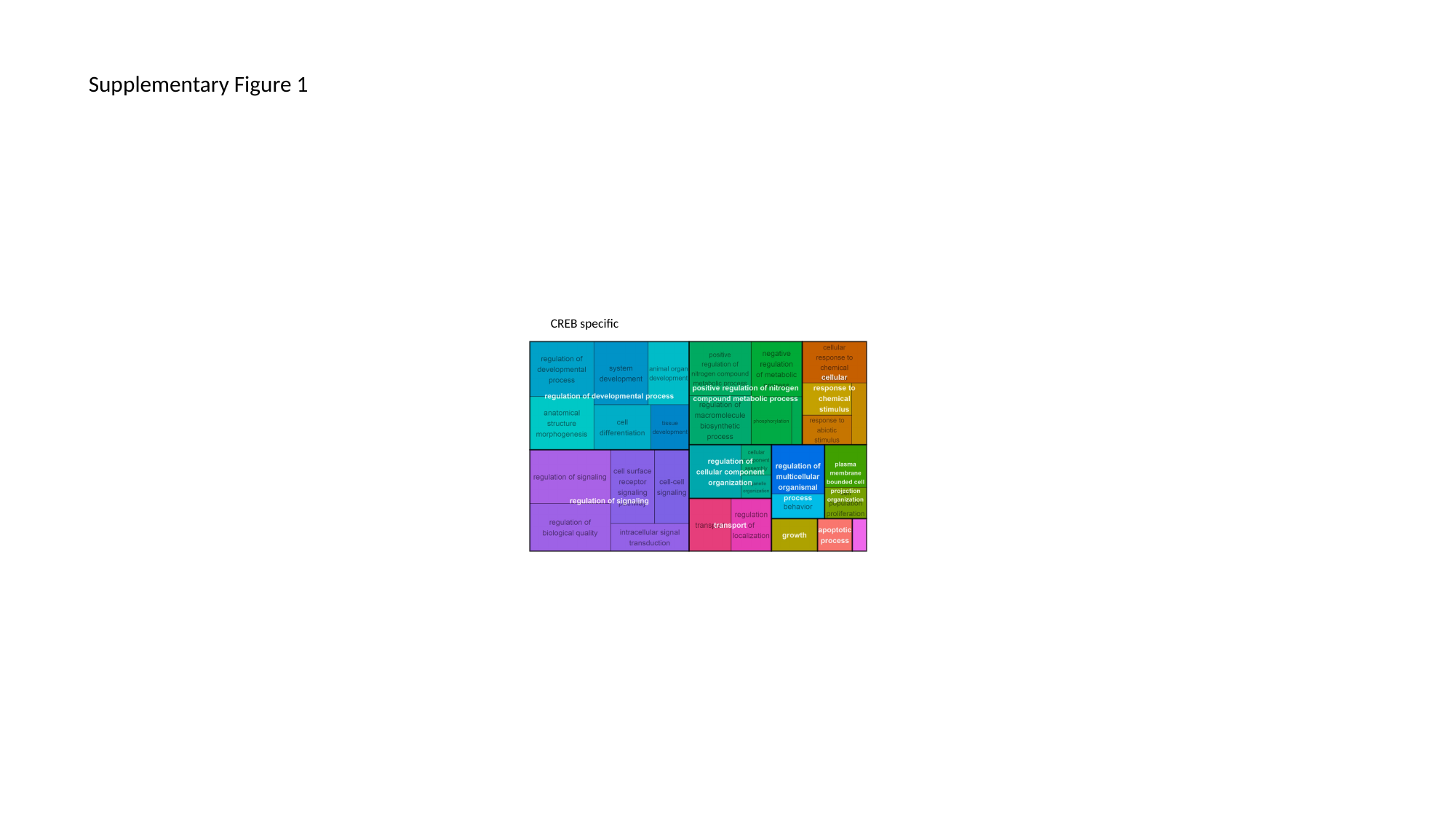

#### Supplementary Figure 1
CREB specific

### Slide 2
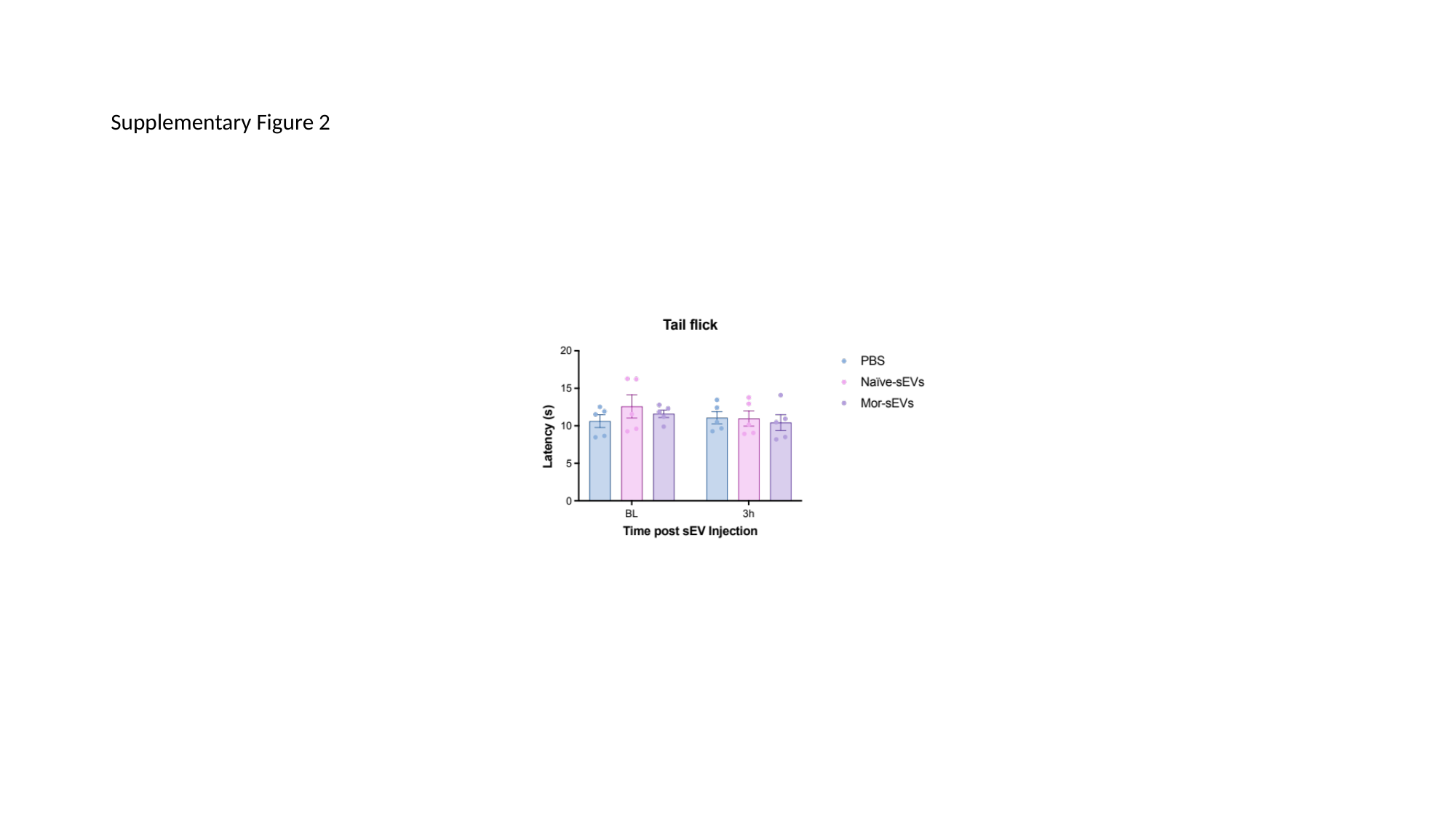

#### Supplementary Figure 2

### Slide 3
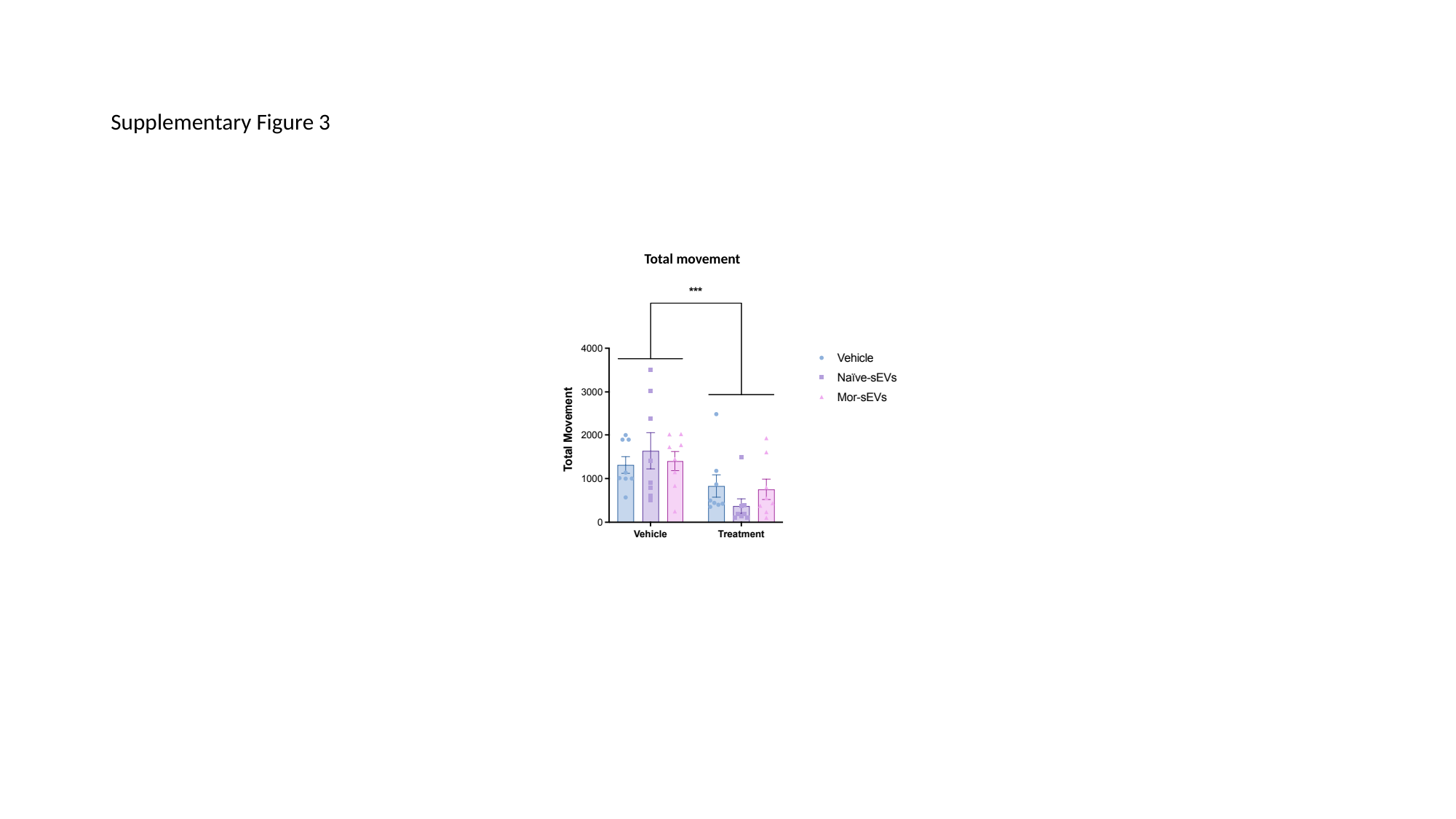

#### Supplementary Figure 3
Total movement
